# Differential antiviral activities of RSV inhibitors in human airway epithelium

**DOI:** 10.1101/250639

**Authors:** Carmen Mirabelli, Martine Jaspers, Mieke Boon, Mark Jorissen, Mohamed Koukni, Dorothee Bardiot, Patrick Chaltin, Arnaud Marchand, Johan Neyts, Dirk Jochmans

**Affiliations:** Laboratory of Virology and Chemotherapy, Department of Microbiology and Immunology, Rega Institute for Medical Research, Katholieke Universiteit Leuven, B-3000 Leuven, Belgium.; Research Group Oto-Rhino-Laryngology, Katholieke Universiteit Leuven and Leuven University Hospitals, B-3000 Leuven, Belgium.; Department of Pediatrics, Pediatric Pulmonology, University Hospital Leuven; Department of Development and Regeneration, Organ Systems, Katholieke Universiteit Leuven; Cistim Leuven vzw, Bioincubator 2, Gaston Geenslaan 2, 3001 Leuven, Belgium; Center for Drug Design and Development (CD3), KU Leuven R&D, Waaistraat 6, 300 Leuven, Belgium

**Keywords:** RSV, antivirals, HuAEC, ciliary beat frerquency, RANTES

## Abstract

We report the use of reconstituted 3D-human airway epithelium cells of bronchial origin (HuAEC) in an air-liquid interface to study respiratory syncytial virus (RSV) infection and to assess the efficacy of RSV inhibitors in (pre-)clinical development. RSV-A replicates efficiently in HuAEC and viral RNA is shed for weeks after infection. RSV infection reduces the ciliary beat frequency of the ciliated cells as of 4 days post infection, with complete ciliary dyskinesia observed by day 10. Treatment with RSV fusion inhibitors resulted in an antiviral effect only when added at the time of infection. In contrast, the use of replication inhibitors (both nucleoside and non-nucleosides) elicited a marked antiviral effect even when start of treatment was delayed until one or even three days after infection. Levels of the inflammation marker RANTES (mRNA) increased ∼200-fold in infected-untreated cultures (at three weeks post infection), but levels were comparable to those of uninfected cultures in the presence of PC-876, a RSV-replication inhibitor, demonstrating that an efficient antiviral treatment inhibits virus induced inflammation in this model. Overall, HuAEC offer a robust and physiologically relevant model to study RSV replication and to assess the efficacy of antiviral compounds.

## INTRODUCTION

The human respiratory syncytial virus (RSV) is worldwide the most prevalent viral pathogen associated with acute lower respiratory infection (ALRI) in infants and children ^1^. Based on data collected in 2015, an estimated 33 million episodes of RSV-ALRI resulted in about 3.2 million hospital admissions and 59 600 in-hospital deaths in children younger than 5 years, of which 27 300 in children younger than 6 months ^2^. RSV causes also significant disease in elderly as well as in immunocompromised and transplant recipients ^3^. Currently, infected patients mainly receive symptomatic treatment and high-risk young paediatric patients (premature, with congenital cardiac abnormality or chronic lung disease) receive prophylactic treatment with the monoclonal antibody Palivizumab ^4^. Fourteen trials with RSV vaccines and vaccine-like monoclonal antibodies (mAbs) are currently ongoing but the development of a safe and effective vaccine for all the ` at risk` populations remains challenging ^5^. Ribavirin (RB V) is currently the only small molecule, which has been approved for treatment of severe RSV infections by aerosol administration, but there is no clear proof of efficacy ^6^.

In recent years, several direct acting RSV-inhibitors have entered clinical development, i.e. fusion inhibitors such as Presatovir (GS-5806) and JNJ-678, and the viral polymerase, nucleoside inhibitor Lumicitabine (ALS-8176) ^7,8^. Significant inhibition of RSV replication in human healthy volunteers experimentally challenged with RSV has been reported ^9^. Fusion inhibitors have a low barrier to resistance development; a single mutation in the viral target protein F compromise their antiviral activity. In addition, different classes of fusion inhibitors are typically cross-resistant ^10^. In contrast, the *in vitro* barrier to resistance of RSV nucleoside polymerase inhibitors [such as ALS-8176], has been shown to be very high and multiple mutations in the active site of the polymerase are required for the virus to acquire a resistant phenotype ^11^. Another promising RSV inhibitor in active pre-clinical development is PC786, a non-nucleoside inhibitor of RSV replication ^12^. Its exact mechanism of action is not fully understood but it is clearly different from ALS-8176 as both classes differ chemically (non-nucleoside vs nucleoside) and no cross-resistance is observed.

The study of RSV antivirals has been standardized *in vitro* in cell lines, such as HEp-2 and HeLa, which permit reproducible assays at sufficient throughput. Here, we explore the use of a fully differentiated human airway epithelium of bronchial origin on an air-liquid-interface setup to assess the efficacy of different classes of RSV inhibitors. This 3D culture system contains all relevant cell types of the lower respiratory tract (ciliated cells, goblet cells, mucus-producing cells) except for cells of the immune system. This system proved valuable to study infections with RSV and other respiratory virus infections ^13^.

## MATERIALS and METHODS

### Media, Cells, Virus and Compounds

Dulbecco’s Modified Eagle Medium with L-glutamine and high glucose and without pyruvate (DMEM cat no 41965-039), Dulbecco’s Phosphate Buffered Saline (PBS cat no 14190-094) and non-essential amino acids solution (NEAA cat no 11140-035) were obtained from Thermo Fisher Scientific. Fetal bovine serum was obtained from Hyclone (cat. no SV30160.03) and heat inactivated at 56°C for 30 minutes. HEp-2 cells and RSV-A Long strain were obtained from ATCC (cat. no CCL-23 and VR-26 respectively). Human airway epithelium of bronchial origin in an air-liquid interface cell culture system (HuAEC) and Mucil Air Medium were obtained from Epithelix, Switzerland (cat. no EP01MD and EP04MM respectively). ALS-8112 and ALS-8176 ^11^, AZ-27 ^14^ and PC786 (patent WO2016/055791) were synthesized according to published procedures. GS-5806 and TMC353121 were obtained from MedChemExpress (NJ, USA) and ribavirin was from ICN Pharmaceuticals (CA, USA).

### Virus and cells for *in vitro* standard antiviral assay

HEp2 cells were seeded at 5.10^3^ cells/well in a 96-well plate and were further cultured in DMEM (supplemented with 2% FBS and 1% NEAA) at 37°C and 5% CO_2_. The day after, medium was replaced by a serial dilution (1:2) of the antiviral test compounds in the same medium and the cultures were infected with RSV-A Long strain at a multiplicity of infection (MOI) of 0.01 CCID_50_/cell. After five days, the typical cytopathic effect (CPE) was scored microscopically in all wells, on a scale of 0-5. The 50% effective concentration (EC_50_) was calculated by logarithmic interpolation as the concentration of compound that results in a 50% protective effect against virus-induced CPE. Potential inhibition of cellular metabolic activity by all the compounds was evaluated in a similar setup in treated-uninfected cultures where metabolic activity was quantified at day 5 by using the MTS readout. The 50% cytotoxic concentration (CC_50_) was calculated by logarithmic interpolation as the concentration of compound that results in a 50% decrease of MTS signal.

### Infection of HuAEC inserts

HuAEC of bronchial origin from healthy donors, were provided in porous culture inserts on an air-liquid interphase setup and were maintained with Mucil Air medium at 37°C and 5% CO_2_ for at least 4 days before use. Each insert contained 4 x 10^5^ cells (epithelix.com). An apical wash with PBS (300 μL) was performed just before infection with RSV-A Long strain (10 CCID_50_/insert). After 2 hours, the inoculum was removed and an apical wash with PBS was collected (T0 of infection). Medium in the basal compartment of the HuAEC (basal medium) was refreshed concomitantly. Apical washes were collected at indicated times p.i. and stored at -20°C until later RNA extractionorat-80°C in culture medium supplemented with 20%-sucrose for long term storage. Antiviral compounds were added to the basal medium and refreshed daily for the duration of the treatment window at the indicated concentrations according to three regimens: (i) prophylactic treatment, the antiviral compound was administered 2 h before infection until day 4 p.i., (ii) early therapeutic treatment, the compound was added from day 1 to day 5 p.i, (iii) late therapeutic treatment, the antiviral compounds was added from day 3 until day 7 p.i. For untreated cultures, the same concentration of DMSO as the drug treated-inserts was added to the culture medium. Each condition tested was performed in duplicate.

### RNA extraction and RT-qPCR

Apical washes collected in PBS were subjected to RNA extraction using the NucleoSpin kit (Macherey-Nagel) according to manufacturer instruction. Viral RNA was quantified by means of RT-qPCR with the iTaq universal One Step Kit (BIORAD) on a Lightcycler 96 (Roche) termocycler. RSV-A primers and probe for amplification were previously described ^15^. Genome copies values were calculated by using a standard curve obtained with increasing dilution of a synthetic GeneBlock (IDT technologies), corresponding to the sequence of the amplicon. Data were plotted with GraphPad software.

### Measurement of ciliary beat frequency (CBF)

At 4, 10 and 15 days p.i., inserts were transferred in a 6 well plate with a small meniscus of medium. An inverted microscope (Leitz, Labovert FS) was used at a magnification of 600X to acquire images by a MotionScope high speed camera at temperature of 22°C. Images (at least 1024 images) were captured at 512 frames/sec in at least three different areas of the insert. The ciliary beat frequency (CBF) value was computed using a custom-made Matlab software previously described ^16,17^. Briefly, the region of interest (ROI) was defined as all pixels with significant motion. Then, CBF values were calculated for each individual pixel in the ROI by Fast Fourier analysis and expressed as a histogram of CBF values for all the pixels in the ROI. The mean CBF value of this histogram was used as result for one CBF measurement. The average of at least three measurements per insert, i.e. six measurements per condition is reported.

### RANTES mRNA levels determination by RT-qPCR

HuAEC were harvested and cellular RNA was extracted with the RNA extraction kit (RNeasy Mini Kit, Qiagen). Specific primers for RANTES were designed and (3-actin was used as an internal control. Samples were quantified by qPCR with the iTaq universal Sybr Green One Step Kit (BIORAD) on a Lightcycler 96 (Roche) thermocycler. Fold change from the uninfected-untreated control was calculated according to the ΔΔCt method.

## RESULTS

### Anti-RSV activity of selected reference antivirals in HEp-2 cells

The *in vitro* anti-RSV activity of a selection of reference compounds was first assessed in a CPE-based antiviral assay in HEp-2 cells in order to determine a common administration dose for the HuAEC system (Table 1). The (i) entry/fusion inhibitors: GS-5806 ^18^ and TMC353121 ^19^; (ii) the nucleoside viral polymerase inhibitor ALS-8112 and its prodrug ALS-8176 ^11^; (iii) two non-nucleoside replication inhibitors AZ-27 ^20^ and PC786 ^12^ as well as (iv) the pan-antiviral ribavirin (RBV) were included in the study. Fusion inhibitors elicited the most potent *in vitro* anti-RSV activity, followed by the non-nucleoside inhibitors and the nucleoside RSV inhibitors. Toxicity was evaluated in parallel and all the direct-acting anti-RSV compounds showed CC_50_ values in a concentration range at least 2- log_10_ higher than the EC_50_, highlighting a good selectivity index. RBV proved the least active compound with a significant cytotoxicity in HEp-2 cells.

**Table 1.** **Antiviral activity of reference compounds in HEp2 cells**

|  | EC<sub>50</sub><sup>a</sup> (nM) |  | CC<sub>50</sub><sup>b</sup> (nM) |  |
| --- | --- | --- | --- | --- |
|  | Median (IQR) | N | Median (IQR) | N |
| **GS-5806** | 0.31 (0.20-0.51) | 12 | > 100 | 14 |
| **TMC353121** | 0.46 (0.28-0.66) | 65 | > 100 | 110 |
| **ALS-8176** | 470 (130-1300) | 10 | > 50000 | 18 |
| **ALS-8112** | 1300 (650-2000) | 25 | > 50000 | 8 |
| **Ribavirin** | 11000 (8900-15000) | 164 | 42000 (33000-47000) | 70 |
| **AZ-27** | 21 (14-39) | 6 | > 50,000 | 9 |
| **PC786** | 1.2 (0.81-1.3) | 11 | > 50000 | 13 |

<sup>a</sup>CPE scoring in RSV/HEp2 cell culture system
<sup>b</sup>Toxicity on HEp2 cells using viability staining with MTS
IQR: Interquartile Range
N: number of replicates

### Fusion inhibitors exert antiviral activity on RSV-infected HuAEC only with a prophylactic regimen

HuAEC inserts were infected with RSV-A (Long strain) and after one day (early treatment) or 3 days (late treatment), the fusion inhibitors GS-5806 and TMC353121 were added to the medium of the basal compartment at a concentration of 100-fold their EC_50_ in HEp-2 cells. RSV RNA levels were quantified in the apical washes at indicated times post infection. No antiviral effect was observed neither with the early treatment (Fig. 1, left panel) nor with late treatment (data not shown). By contrast, when GS-5806 (1000-fold *in vitro* EC_50_) and TMC353121 (100-fold *in vitro* EC_50_) were added 2 h before infection, a significant reduction in viral RNA levels was detected (>2 log_10_) by day 4 p.i. (Fig. 1, right panel). However, upon removal of the inhibitor from the culture medium, viral rebound was observed.

**Fig. 1.**
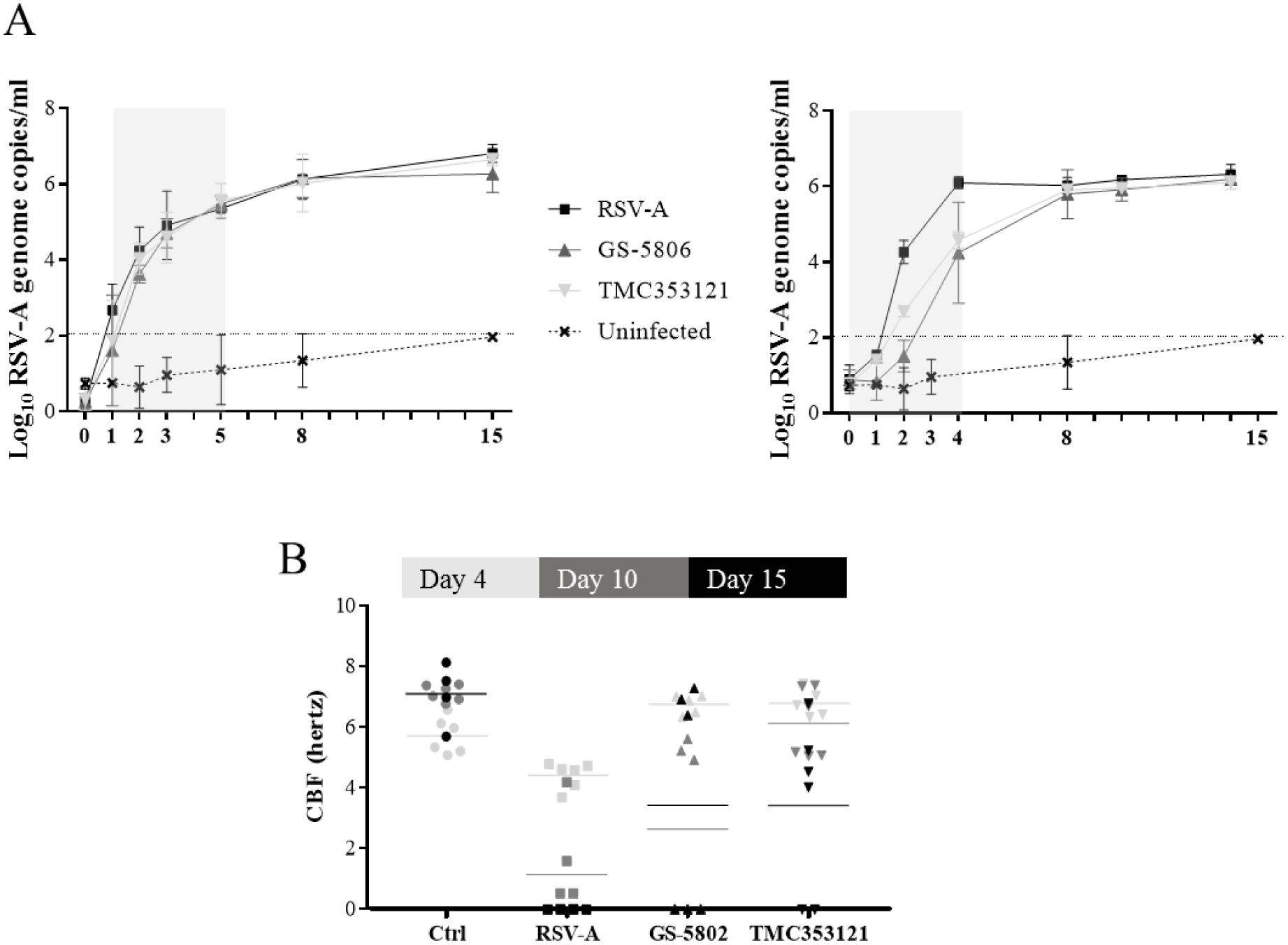
The effect of the RSV fusion inhibitors GS-5806 and TMC353121 was evaluated in HuAEC by using a therapeutic-early treatment regimen (day 1-5 p.i.) (left panel) or by using a prophylactic regimen (-2 h to day 4 p.i) (right panel). Compounds were added to the basal medium at a concentration of 100-fold their *in vitro* EC_50_. Apical washes were collected at selected time points p.i. and viral RNA was quantified by means of RT-qPCR. The treatment period is depicted in grey. The sensitivity threshold was determined by the highest RSV RNA signal quantified on uninfected-untreated cells (ctrl). Error bars represent the standard deviation of duplicates. Ciliary beat frequency was measured by high-speed video-microscopy with a custom-made Mathlab script. Three areas per inserts, i.e. six measurements per condition were acquired at three data points: 4-10 and 15 days p.i. Average of the measurements per condition are represents as bars (B).

Infection of the cultures with RSV did not results in any noticeable cytopathic effect, therefore the quantification of the ciliary beat frequency (CBF) was used (in addition to quantifying of viral genomes levels) to assess the impact of the compound-treatment on viral infection. Ciliary motility was quantified for at least three different areas per insert, i.e. at least six measurements per condition (Fig. 1B). In line with an earlier report [20], a slight change in CBF was observed at day 4 p.i., with 4.41 +/-0.39 Hz in infected-untreated versus 5.71+/-0.54 Hz in uninfected inserts. By day 10 p.i, all the analyzed areas of RSV-infected untreated cultures showed complete dyskinesia (CBF: 1.1 +/- 1.4 Hz). In infected cultures, the treatment with fusion inhibitors delayed ciliary dyskinesia and in some of the areas of the drug-treated HuAEC cultures a CBF of 5-6 Hz was still noted.

### Polymerase inhibitors, nucleoside and non-nucleoside, show anti-RSV activity in a therapeutic setup

The potential anti-RSV activity of ALS-8112 and its prodrug ALS-8176 (both at 100-fold their EC_50_) was assessed in HuAEC with an early (day 1 to day 5 p.i.) and late therapeutic treatment (day 3 to day 7). The early treatment resulted in a drop of viral RNA levels to (or lower than) the levels of the uninfected-untreated control (sensitivity threshold) and remained nearly undetectable, with a 4- and 6- log_10_ yield reduction by the end of the experiment (day 15 p.i.) for ALS-8176 and ALS-8112, respectively (Fig. 2A, left panel). When the compounds were first added to the inserts at day 3 p.i. ‐when viral RNA levels had reached the plateau phase‐ a marked reduction (∼2-log_10_) in viral RNA was observed by day 7 p.i. After the end of treatment, no viral rebound was detected (Fig. 2A, right panel). The prodrug ALS-8176 was slightly less potent than the parental compound.

**Fig. 2.**
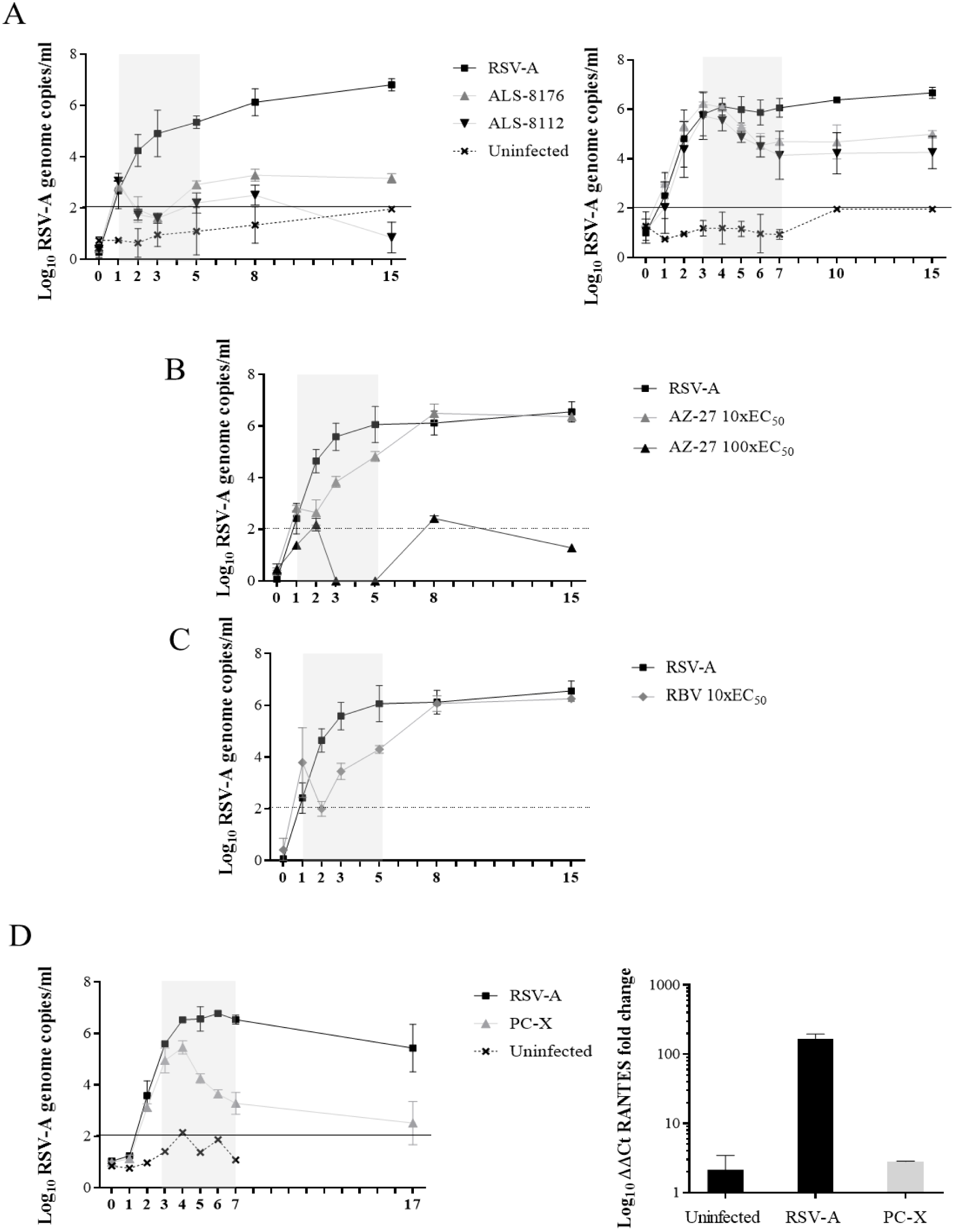
HuAEC were infected with RSV-A (Long strain) and were treated with (i) the nucleoside viral polymerase inhibitor ALS-8112 and prodrug ALS-8176 (A) at 100-fold the *in vitro* EC_50_ by using either a therapeutic-late (day 3-7 p.i., left panel) or early (day 1-5 p.i., right panel) treatment regimen or (ii) the replication inhibitor AZ-27 (B) at 10-fold and 100-fold the *in vitro* EC_50_ or with ribavirin (C) at 10-fold EC_50_ with a therapeutic-early treatment schedule or (iii) with PC876 (D, left panel) at 100-fold the *in vitro* EC_50_ by using a therapeutic-late treatment schedule. The treatment period is depicted in grey. The sensitivity threshold was determined by the highest RSV-RNA signal quantified on uninfected-untreated cells (ctrl). Error bars represent the standard deviation of duplicates. HuAEC were harvested at day 24 p.i. and total cellular mRNA was extracted. Levels of RANTES mRNA were quantified by means of RT-qPCR (D, right panel). β-actin was used as internal control. Fold change of the untreated-uninfected control was calculated with the ΔΔCt method. Error bars represent the median of three independent quantifications of the condition and each condition was performed in duplicate (i.e. six measurement).

Next, the potential anti-RSV activity of the non-nucleoside replication inhibitor AZ-27 was assessed with an early therapeutic treatment regimen at concentrations of 10-fold and 100-fold the *in vitro* EC_50_. At 10-fold the EC_50_, a rapid decrease of viral RNA was observed after start of treatment, followed by an equally rapid viral rebound after cessation of treatment (Fig. 2B). To study whether the rebound resulted from the emergence of drug-resistant variants, the viruses from the rebound phase were sequenced. However, no mutations were detected in the AZ-27-target (L-) gene. At 100-fold the *in vitro* EC_50_, the effect of AZ-27 proved comparable to that of ALS-8112, with no detectable viral RNA on days 1-2 after start of treatment and no viral rebound after stop of treatment (Fig. 2B). Interestingly, akin AZ-27 treatment at 10-fold the EC_50_, RBV at 10-fold the EC_50_ resulted also in a transient, sub-optimal antiviral effect (Fig. 2C).

Another non-nucleoside replication inhibitor, PC786 (at 100-fold the EC_50_) resulted in the most pronounced antiviral activity with a rapid decline of viral RNA yield in the apical washes even if the compound was first added to the inserts 3 days after infection. At the end of treatment, (at day 7 p.i.), a 3-log_10_ reduction in viral RNA was noted and no rebound of viral replication was observed (Fig. 2D, left panel). To explore whether antiviral treatment affected virus-induced immune activation of the HuAEC system; mRNA levels of the cytokine RANTES were quantified. This cytokine has been reported to be highly induced following RSV infection ^21^. RANTES mRNA levels were ∼200x higher in infected-untreated *versus* uninfected cultures (Fig. 2D, right panel). Notably, in PC786-treated infected cultures, RANTES mRNA levels were comparable to those of the uninfected-untreated controls.

## DISCUSSION

We present a model of reconstituted human airway epithelium of bronchial origin to study the efficacy of small molecules inhibitors of RSV infection in a physiologically relevant context. To validate this model for antiviral studies, we explored the effect of a panel of RSV inhibitors in (pre-)clinical development targeting either viral fusion/entry (TMC35112 and GS-5806) or viral replication (the nucleoside ALS-8176 and the non-nucleosides AZ-27 and PC786).

For fusion inhibitors, an antiviral effect was only noted when the compounds were first added before the infection. Under these conditions, viral kinetics suffered a delay in presence of GS-5806 and TMC353121, regardless of the tested concentration (100 or 1000-fold the *in vitro* EC_50_, respectively) but viral rebound was detected after stop of treatment. Possibly, entry inhibitors prevent only the first round of infection but once the virus has entered the cell and spreads from cell to cell, this class of compounds is no longer able to exert an antiviral effect. In a very recent study, the antiviral effect of another potent fusion inhibitor JNJ-53718678, was explored in HuAEC ^22^. Akin to our study, the compound proved active when added prophylactically (1 h before infection). However, in apparent contrast with our findings, some (weak) activity of JNJ-fusion inhibitor was also detected in a therapeutic regimen (when the compound was added 1-day p.i.). Notably, the JNJ-fusion inhibitor was added to both the basal and the apical compartment of the HuAEC inserts and the experimental endpoint was set at day 4p.i. (the day of stop of treatment). In our study, the inhibitor was only added to the basal compartment and viral replication was monitored daily until day 15 post infection (i.e. 11 days after stop of treatment). The administration of the inhibitor to the basal compartment only, might have affected the efficacy (since the infection occurs in the apical compartment) but might better reflect the situation in the infected patient receiving systemic dosing of the drug. It is noteworthy that in the first GS-5806-clinical trial on 37 healthy adults with low serum RNA-RSV, the compound was administered orally following RSV detection but before symptoms development ^23^. A recent mathematical modelling of the effect of fusion inhibitors on RSV treatment also suggests, in line with our findings, that the activity of fusion inhibitors strongly depends on early treatment ^24^. Despite the lack of an obvious clearance of RSV RNA, treatment with fusion inhibitors delayed the onset of ciliary dyskinesia. By means of high-speed video-microscopy, complete ciliary diskynesia was observed by day 10 p.i. in RSV-infected HuAEC, whereas still some ciliary beat activity in GS-5806- and TMC353121-treated samples was noted at day 15 p.i. Hence, treatment with fusion inhibitors, even sub-optimal, may delay the onset of symptoms and be beneficial in natural infections, particularly for high-risk patients.

ALS-8112 is a cytidine nucleoside analogue and its prodrug, ALS-8176, is currently being evaluated in clinical trials (clinicaltrial.gov). In a human RSV-challenge experiment, ALS-8176 reduced the absolute RSV viral load to undetectable levels in 1.3-2.3 days ^7^. In the HuAEC system, the compound was tested in an early-therapeutic treatment and resulted in a marked antiviral effect. Similarly to the dynamics of activity observed *in vivo,* the virus became undetectable between day 1 and 2 p.i. Also in late therapeutic treatment schedule, a significant antiviral effect was observed.

Overall, treatment with non-nucleoside replication inhibitors was effective both when given early or late after infection; at least when concentrations ≥ 100-fold the *in vitro* EC_50_ were used. Indeed, lower concentrations (10fold i*n vitro* EC_50_) resulted in little effect and in an immediate rebound of viral replication after stop of treatment. Interestingly, no drug-resistant variants were selected following sub-optimal dosing of AZ-27. Among all the inhibitors studied, PC876 elicited the most potent activity in HuAEC model; in particular it still resulted in a profound antiviral effect when first added to the infected cultures three days post infection. In clinical practice early treatment is very difficult to realize, as patients most often seek medical advice 3-4 days after infection. Therefore, drugs that are able to curb viral replication when added late after infection may likely hold most promise as therapeutic agents. Of clinical interest, RBV, the only antiviral approved for the treatment of RSV in high-risk populations but debated in terms of efficacy, proved inefficient to control RSV infection after stop of treatment in our model. Notably, in the clinic, the recommended RBV treatment is oral whereas in our model, we mimicked a system treatment. It remains to address whether an apical administration of RBV in HuAEC will result in a persistent control of viral replication.

An added value of the HuAEC model is the possibility to study the inflammatory response of a differentiated epithelium to viral infection. By monitoring inflammation markers, the efficacy of an antiviral treatment in curing the system and restoring its innate immune state can be studied. We monitored (mRNA levels of) RANTES, a cytokine highly expressed during RSV infection, and noted a ∼200x induction upon infection. Treatment with PC876 completely abrogated this increase; indicating that RANTES is a good marker of RSV infection in HuAEC and that effective antiviral treatment restores the innate immune state of the system.

In conclusion, we describe a robust and reproducible model to study RSV infection and to assess the potency and the antiviral dynamics of RSV inhibitors. 3D *ex-vivo* cultures are largely debated for their clinical relevance as surrogates of human infection. We demonstrate that RSV-infected 3D cultures of human airway epithelium recapitulate aspects of the response to antiviral therapy and provide further evidences that HuAEC are relevant cell-based model for disease and antiviral studies.

## Acknowledgements.

We thank Tina Van Buyten, Kim Donckers and Jasmine Paulissen for excellent technical assistance.

## Transparency declaration.

All the authors have nothing to declare.

